# Increased neuronal expression of the early endosomal adaptor APPL1 replicates Alzheimer’s Disease-related endosomal and synaptic dysfunction with cholinergic neurodegeneration

**DOI:** 10.1101/2024.09.19.613736

**Authors:** Ying Jiang, Kuldeep Sachdeva, Chris N. Goulbourne, Martin J. Berg, James Peddy, Philip H. Stavrides, Anna Pensalfini, Monika Pawlik, Sandeep Malampati, Lauren Whyte, Balapal S. Basavarajappa, Subbanna Shivakumar, Cynthia Bleiwas, John F. Smiley, Paul M. Mathews, Ralph A. Nixon

**Affiliations:** Center for Dementia Research, Nathan S. Kline Institute for Psychiatric Research, Orangeburg, NY 10962, USA; Department of Psychiatry, New York University Grossman School of Medicine, New York, NY 10016, USA; Emotion Brain Institute, Nathan S. Kline Institute for Psychiatric Research, Orangeburg, NY 10962, USA; Department of Cell Biology, New York University Grossman School of Medicine, New York, NY 10016, USA; Neuroscience Institute, New York University Grossman School of Medicine, New York, NY 10016, USA

## Abstract

Endosomal system dysfunction within neurons is a prominent early feature of Alzheimer’s disease (AD) pathology. Multiple AD risk factors are regulators of endocytosis and are known to cause hyper-activity of the early-endosome small GTPase rab5, resulting in neuronal endosomal pathway disruption and cholinergic neurodegeneration. Adaptor protein containing Pleckstrin homology domain, Phosphotyrosine binding domain, Leucine zipper motif (APPL1), an important rab5 effector protein and signaling molecule, has been shown *in vitro* to interface between endosomal and neuronal dysfunction through a rab5-activating interaction with the BACE1-generated C-terminal fragment of amyloid precursor protein (APP-βCTF), a pathogenic APP fragment generated within endosomal compartments. To understand the contribution of APPL1 to AD-related endosomal dysfunction *in vivo*, we generated a transgenic mouse model over-expressing human APPL1 within neurons (Thy1-APPL1 mice). Strongly supporting the important endosomal regulatory roles of APPL1 and their relevance to AD etiology, Thy1-APPL1 mice develop enlarged neuronal early endosomes and increased synaptic endocytosis due to increased rab5 activation. We demonstrated pathophysiological consequences of APPL1 overexpression, including functional changes in hippocampal long-term potentiation (LTP) and long-term depression (LTD), degeneration of large projection cholinergic neurons of the basal forebrain, and impaired hippocampal-dependent memory. Our evidence shows that neuronal APPL1 elevation modeling its functional increase in the AD brain induces a cascade of AD-related pathological effects within neurons, including early endosome anomalies, synaptic dysfunction, and selective neurodegeneration. Our *in vivo* model highlights the contributions of APPL1 to the pathobiology and neuronal consequences of early endosomal pathway disruption and its potential value as a therapeutic target.

**Significance Statement:** Neuronal endosome dysfunction appears early in Alzheimer’s disease (AD) and is linked to memory loss. Genes and risk factors associated with AD often increase rab5 activity, a protein that disrupts endosomal signalling when hyperactivated. APPL1, a key rab5 partner, worsens this dysfunction via its interaction with APP-βCTF, a protein fragment associated with AD. To explore APPL1’s role, we created a genetically modified mouse that overexpresses APPL1 in neurons. This model provides the first *in vivo* evidence that APPL1 overexpression triggers key AD-like effects: rab5 hyperactivation, enlarged early endosomes, loss of cholinergic neurons, reduced synaptic plasticity in memory-related brain regions, and memory deficits. These findings highlight APPL1’s role in AD pathogenesis and its potential as a therapeutic target.

## 1. Introduction

Endosomal pathway dysfunction is the earliest known cytopathology of Alzheimer’s disease (AD), with endosomal anomalies emerging years, possibly decades, before the buildup of hallmark pathologies like β-amyloid and tau tangles (Cataldo et al., 2000; Colacurcio et al., 2018; Pensalfini et al., 2020; Fahimi et al., 2021). Research has shown that individuals carrying the apolipoprotein E (apoE) e4 allele, the major genetic risk factor for late onset AD, develop endosomal alterations in neurons (Cataldo et al., 2000). Similar changes were seen in apoE e4 mouse models without β-amyloid or tau pathology (Nuriel et al., 2017; Peng et al., 2024). Trisomy 21 Down syndrome (DS), which leads to AD in later life, is associated with early endosomes disruption and early emerging cholinergic neurodegeneration in the basal forebrain due to intracellular accumulation of APP-βCTF, a C-terminal fragment of amyloid precursor protein (APP) (Jiang et al., 2016; Kim et al., 2016; Nixon et al., 2017; Colacurcio et al., 2018; Lauritzen et al., 2019). APP-βCTF levels were elevated in human AD brain (Pera et al., 2013; Kim et al., 2016). Across various AD models, higher levels of APP-βCTF contribute directly to early endosome dysfunction and disruptions of endosomal trafficking leading to neuronal loss (Jiang et al., 2010; Jiang et al., 2016; Lauritzen et al., 2016; Nixon et al., 2017; Bourgeois et al., 2018; Colacurcio et al., 2018).

Both APP-βCTF and rab5 proteins on endosome membrane bind to APPL1 (Adaptor Protein containing Pleckstrin homology domain, Phosphotyrosine binding domain, and Leucine zipper motif) via the phosphotyrosine binding domain and pleckstrin homology domain respectively, interactions that lead to the accumulation of APPL1 at rab5-containing endosomal membranes (Miaczynska et al., 2004; Mao et al., 2006; Zhu et al., 2007; Kim et al., 2016; Diggins and Webb, 2017; Xu et al., 2018; Hiragi et al., 2022). APPL1 stabilizes the GTP-bound, endocytic-active form of rab5, which leads to accelerated endocytosis, early endosome enlargement, and impaired transport of endosomes in primary cortical neurons (Kim et al., 2016). Given that directly hyper-activating rab5 *in vivo* in mice via modest neuronal overexpression of rab5 recapitulates AD-related early endosome anomalies as well as their neurodegenerative consequences (Kim et al., 2016; Pensalfini et al., 2020), we undertook this study to determine whether APPL1 stabilization of active rab5 lead to similar early endosomal pathologies, highlighting the role of this regulatory and linking molecule in AD early endosome pathobiology.

APPL1 interacting with rab5 at the early endosomal membrane modulates endocytic trafficking while interfacing with an array of structural and signaling proteins (Miaczynska et al., 2004; Mao et al., 2006; Zhu et al., 2007; Diggins and Webb, 2017; Hiragi et al., 2022). Notably, APPL1 modulates Akt signaling (Majumdar et al., 2011; Bohdanowicz et al., 2012; Wang et al., 2016; Goto-Silva et al., 2019) and mediates nerve growth factor signaling through TrkA (Lin et al., 2006) to support cholinergic neuron function. Following rab5-GTP hydrolysis, APPL1 is released from early endosomal membranes enabling its translocation into the nucleus to regulate gene expression (Rashid et al., 2009; Zoncu et al., 2009). APPL1’s endosomal roles are also important to hippocampal synaptic plasticity (Formolo et al., 2022) including controlling the N-methyl-d-aspartate receptor (NMDAR)-dependent potentiation that facilitates extinction memory retrieval (Hua et al., 2023). Notably, APPL1 levels are abnormally elevated 2-fold in AD brain compared to control (Johnson et al., 2022).

To examine *in vivo* the interface between APPL1 function and the diverse endosomal and neurodegenerative consequences attributed to early endosome pathobiology in AD, we generated a novel transgenic mouse model that overexpresses APPL1 in neurons (Thy1-APPL1 mice). With modest APPL1 overexpression (2-fold), this Thy1-APPL1 model replicates the AD-related endosome phenotype mediated by rab5 hyper-activation and its downstream pathological consequences on neuronal function, including those at hippocampal synapses. Importantly, the number of choline acetyltransferase (ChAT)-positive neurons in the medial septal nucleus (MSN) was reduced in Thy1-APPL1 mice. Hippocampal-dependent memory, measured by the novel object recognition test, was impaired in Thy1-APPL1 mice. These *in vivo* findings support the idea that APPL1 recruitment to early endosomes is sufficient to replicate endosomal, functional, and neurodegenerative changes induced by elevated APP-βCTF levels and rab5 over-activation in AD and related animal models (Jiang et al., 2016; Nixon et al., 2017; Pensalfini et al., 2020; Jiang et al., 2022; Peng et al., 2024).

## 2. Methods

### 2.1. Thy1-APPL1 transgene construct, generation of transgenic mice, and genotyping

Transgenic mice overexpressing human APPL1 (NM_012096) were generated by Ingenious Targeting Laboratory (Ronkonkoma, NY) using the pTSC21 vector consisting of a murine Thy1 expression cassette (Pensalfini et al., 2020) and a Flag-tag for identification. Two founder mice (M1 and F2) were obtained on a mixed background of 129SvEC and C57BL/6, which were then crossed with C57BL/6 mice (JAX stock number 000664, The Jackson Laboratory, Bar Harbor, ME) for over 10 generations to establish the line used in this study; offsprings from founder F2 mice were used for the current study. Thy1-APPL1 mice genotyping was accomplished by PCR using the following primer pair: forward-5’ ATGAAGTCACCCAGCAGGGAG and reverse 5’ AGGTCAGGTGTGTTGCTGCAC. Each reaction consisted of H_2_O (23 µl), 2x GO Taq master mix (M7122, Promega), primer pair (0.5 µl each), and tail DNA (2 µl of a 0.05-0.1 mg/ml solution). The PCR conditions were as follows: 94°C for 3 minutes, followed by 30 cycles of 94°C for 30 seconds, 58°C for 30 seconds, and 72°C for 30 seconds, with a final extension step at 72°C for 5 minutes. The reaction mixes were then loaded onto agarose gels, Thy1-APPL1 mice were positively identified by a band presented at or above 500 bp. Thy1-APPL1 mice bred normally with the expected Mendelian ratio of offsprings, with normal body weight and lifespan. Throughout, non-transgenic (non-Tg) wild-type littermate mice were used as controls.

Mouse experimentation and animal care were approved by the Institutional Animal Care and Use Committee (IACUC) of the Nathan S. Kline Institute. The approval is renewed every three years, and the current protocol number is AP2024-746. Mice were housed in the NKI Animal facility and kept under a 12-hour day and night cycle, at temperatures of approximately 70°F (±2°C) and humidity between 40% and 60%. Both male and female mice were used.

### 2.2. Immunocytochemistry and Western blot analysis

Thy1-APPL1 mice and age-matched non-Tg were anesthetized with a combination of ketamine (100 mg/kg BW) and xylazine (10 mg/kg BW) and transcardially perfused with 0.9% NaCl in 0.1 M phosphate-buffered (pH 7.4) as described earlier (Yang et al., 2014). One hemibrain was frozen and stored at -80^0^C for biochemical analysis (see below), while the other hemibrain was drop-fixed in 4% paraformaldehyde (PFA) in 0.1M sodium cacodylate buffer, pH 7.4 (Electron Microscopy Sciences EMS) for immunolabeling as previously described (Pensalfini et al., 2020; Jiang et al., 2022). After more than 48 hours of fixation, 40-μm thick vibratome brain tissue sections from various brain regions were collected and washed three times with antibody dilution buffer containing bovine serum albumin (BSA; 1%; Sigma), saponin (0.05%; Sigma), and normal horse serum (NHS; 1%; Thermo Fisher) in Tris-buffered saline (TBS, pH 7.4), then blocked with 20% NHS in TBS for one hour at room temperature before incubation with commercial antibodies against rab5 (Abcam Cat# ab18211, RRID:AB_470264; 1:500) (Kim et al., 2016; Pensalfini et al., 2020), rab5-GTP (active rab5: NewEast Biosciences Cat# 26911, RRID:AB_2617182; 1:50) (Kim et al., 2016; Pensalfini et al., 2020), ChAT (Millipore Cat# AB144P, RRID:AB_2079751; 1:250) (Jiang et al., 2016; Pensalfini et al., 2020), APPL1 (Proteintech Cat# 12639-1-AP, RRID:AB_2289669; 1:500) (Kim et al., 2016), and the Flag-tag (Thermo Fisher Scientific Cat# MA1-91878-D488, RRID:AB_2537621; 1:500), NeuN (Millipore, Cat# MAB377, RRID:AB_2298772; 1:2000) (Lee et al., 2018), either alone or in combination and visualized with either biotinylated secondary antibodies (Vector Laboratories; Cat# BA-9200, BA-1000 and BA-6000; 1:500) or fluorescence-conjugated secondary antibodies (Thermo Fisher; Cat# A10037, A21206 and A21099; 1:500), as described in our previous studies (Pensalfini et al., 2020; Jiang et al., 2022). An additional mouse monoclonal antibody against APPL1 (Novus Cat#NBP2-46536; RRID:AB_3083472; 1:500) was used for co-immunolabeling with a rabbit polyclonal rab5 antibody (Abcam Cat# ab218624). Confocal images were collected using a Zeiss LSM880 laser confocal microscope and Zen 2.1-Sp3 software. Rab5 and rab5-GTP positive puncta, including intensity, number, average size, and total area per cell, were determined by Fiji/ImageJ 2.3.0 (https://imagej.net/Fiji). Three or more mice per genotype were analyzed, and 20-30 neurons from each mouse were quantified as previously reported (Pensalfini et al., 2020; Jiang et al., 2022). For protein analyses, mouse hemi-brains were homogenized in 1:10 (brain weight: buffer) pH 7.4 buffer containing 250 mM sucrose, 20mM Tris-HCl, 1mM EDTA, 1mM EGTA, protease/phosphatase inhibitors (Pensalfini et al., 2020; Jiang et al., 2022), and western blot analyses as described previously were performed with antibodies against APP (C1/6.1; 1µg/ml) (Jiang et al., 2010), APP-βCTF (M3.2; 2µg/ml) (Morales-Corraliza et al., 2009), APPL1 (Proteintech; 1:1000), rab5 (Abcam; 1:2000), synaptophysin (Sigma-Aldrich Cat# S5768, RRID:AB_477523; 1:2000) (Pensalfini et al., 2020), phosphorylated tau (PHF1; 1:1000, kind gift of P. Davies (Pensalfini et al., 2020) and total tau protein (Agilent Cat# A0024, RRID:AB_10013724; 1:5000) (Pensalfini et al., 2020). All the secondary antibodies for western blot analyses were used according to the manufacturer’s recommendations (Jackson ImmunoResearch Laboratories, PA). As an internal loading control, total protein on each blot was determined by Revert^TM^ 700 Total protein Stain (Licor; Cat# 926-11021) (Wijewantha et al., 2024). Digital gel imager (Syn-gene G:Box XX9) was used to capture both the total protein stain and the ECL images from each blot. Band intensities on each blot were quantified with a combination of Fiji/ImageJ 2.3.0. and Multi Gauge (Fujifilm; V3.0) software.

### 2.3. ELISA of Aβ40 and Aβ42 of DEA extracts from brain tissue of Thy1-APPL1 and non-Tg mice

Aβ40 and Aβ42 levels were determined by sandwich ELISA according to previously published protocols (Schmidt et al., 2012; Jiang et al., 2016). Briefly, brain homogenates were extracted with 0.4% diethylamine (Sigma, Cat# D3131) /100 mM NaCl solution (1:1 of v/v), then processed further with a tissue grinder before centrifugation at 43,000 rpm for one hour at 4°C in a Beckman Coulter TLA 100.3 rotor. The collected supernatants were mixed with 10% volume of 0.5 M Tris buffer (pH 6.8), 100 µl of the buffered samples were then loaded into ELISA plates coated with specific anti-Aβ40 or Aβ42 antibodies as reported previously (Schmidt et al., 2005; Jiang et al., 2016). Horseradish peroxidase (HRP) conjugated secondary antibody (M3.2, 1:1000) (Choi et al., 2009; Jiang et al., 2016) was used for both Aβ40 and Aβ42 detections, and a multimode plate reader (SpectraMax M5, Molecular Devices) was used to read the optical density (O.D.) at 450 nm. The level of Aβ40 or Aβ42 was presented in fmol/g of brain tissue, calculated according to a standard curve using synthetic mouse Aβ40 or Aβ42 peptides (American Peptide Co.) (Choi et al., 2009; Jiang et al., 2016; Pensalfini et al., 2020).

### 2.4. Stereological counting of medial septal nucleus ChAT-immunoreactive neurons

For ChAT immunolabeling, every third consecutive vibratome brain tissue section containing medial septal nucleus (MSN) was treated with 3% H_2_O_2_ before blocking with 20% NHS in TBS for one hour at room temperature, followed by incubation with anti-ChAT antibody (Millipore Sigma; 1:250) as described previously (Jiang et al., 2016; Jiang et al., 2022). Visualization was achieved using diaminobenzidine (DAB; Cat# SK-4100; Vector Laboratories) after incubation with a biotinylated secondary antibody (1:500; Vector Laboratories) and the Vectastain ABC kit (Vector Laboratories Cat# PK6105). To quantify ChAT-positive neurons in the MSN, the optical fractionator method was used combined with ImageJ software as previously described (West et al., 1991; Smiley et al., 2012; Jiang et al., 2016; Jiang et al., 2022). Cell counts were corrected for the z-axis section thickness, which was measured three times per section. The dissector size and density were optimized to maintain a coefficient of error below 0.1.

### 2.5. Electrophysiology

Thy1-APPL1 and non-Tg littermate mice (n=5 for each genotype) were used to measure long-term potentiation (LTP) and long-term depression (LTD) in the hippocampal CA1 as detailed in previous studies (Subbanna et al., 2013; Pensalfini et al., 2020). Mice were sacrificed by cervical dislocation and the hippocampus were immediately collected, followed by sectioning to 400 µm slices. The hippocampal slices were then placed into a recoding chamber filled with artificial cerebrospinal fluid (124 mM NaCl, 4.4 mM KCl, 1.0 mM Na_2_HPO_4_, 25 mM NaHCO_3_, 2 mM CaCl_2_, 2 mM MgSO4, 10 mM glucose, osmolarity between 290-300 Osm/L) at 29°C with 95% O_2_ and 5% CO_2_. CA1 field excitatory postsynaptic potentials (fEPSPs) were recorded by placing the stimulating and recording electrodes in the hippocampal CA1 stratum radiatum. LTP was induced by theta-burst stimulation (TBS, 4 pulses at 100 Hz, with the bursts repeated at 5 Hz, and each tetanus including three 10-burst trains separated by 15 seconds) and LTP responses were recorded for 2 hours. LTD was induced by low-frequency stimulation (LFS, 1 Hz for 900 seconds) and LTD responses were recorded for 80 minutes. For both LTP and LTD experiments, a 10-minutes baseline was recorded at 1-minute intervals with stimulus to evoke a response about 35% of the maximum evoked response. Both LTP and LTD fEPSP slopes were expressed as percentage of baseline (the average value 10 mins before stimulation). The results are expressed as both fEPSP slopes and combined bar graphs of the averages of fEPSP slopes at indicated time points (Mean ± SEM) with individual mouse data as previously reported (Subbanna et al., 2013; Pensalfini et al., 2020).

### 2.6. Electron Microscopy and post-embedding immunogold EM

Electron microscopy and post-embedding immunogold EM were performed on brain sections as previously described (Yang et al., 2009; Pensalfini et al., 2020). Mice were transcardially perfused with a solution of 4% PFA and 2% glutaraldehyde in 0.1 M sodium cacodylate buffer, pH 7.4. Fixed tissue was cut into 80 μm-thick sagittal vibratome sections and post-fixed in 1% osmium tetroxide. Following alcohol dehydration, tissue sections were infiltrated with increasing concentrations of Spurr resin and then embedded flat in Aclar sheets. Regions of interest for ultrastructural analyses, including the prefrontal cortex and hippocampal CA1, were excised, and 50 nm ultrathin sections were prepared and stained with uranyl acetate and lead citrate. The thin sections were then viewed using a Thermo Fisher Talos L120C transmission electron microscope operating at 120 kV. For endosome quantification, approximately 60 EM images (17,500X) per mouse were acquired. The images contained dendritic and synaptic profiles in the proximity (within 5-10 µm) of the neuronal soma located within the pyramidal cell layer V of the pre-frontal cortex (n=7 for both non-Tg and Thy1-APPL1 mice) and CA1 of the hippocampus (n=4 for both genotypes). The average size and circumference of endosomes for each mouse genotype were determined using Fiji/ImageJ (Schindelin et al., 2012). For immunogold EM, ultrathin tissue sections were mounted on nickel grids, air-dried, and then etched for 5 minutes with 1% sodium metaperiodate in PBS. After washing with filtered double-distilled water, the ultrathin tissue sections were incubated with 1% BSA in PBS for 2 hours, followed by overnight incubation with the combination of mouse anti-APPL1 antibody (dilution of 1:2) and rabbit anti-Rab5 antibody (dilution of 1:2) in a humidified chamber at 4 °C. These sections were incubated with gold-conjugated secondary anti-mouse (size 10 nm) and anti-rabbit (size 6 nm) antibodies for 2 hours at room temperature before imaging. For colocalization quantification, approximately 40 EM images (22,000X and 57,000X) per mouse were acquired from the pyramidal cell layer V of the pre-frontal cortex and CA1 of the hippocampus of non-Tg and Thy1-APPL1 mice (n=3 each genotype). Rab5-immunoreactive endosomes, as well as rab5/APPL1 dual-immunoreactive endosomes, were counted manually, with results presented as percentage of rab5^+^/APPL1^+^endosomes versus total rab5^+^ endosomes.

### 2.7. Synaptosome isolation

Synaptosomes were isolated from non-Tg and Thy1-APPL1 mouse hippocampi (n=6 mice per genotype). For hippocampal synaptosomes, hippocampi were homogenized in 250 µl homogenization buffer (0.32 mol/L sucrose, 0.1 mmol/L CaCl_2_, 1 mmol/L MgCl_2_ with protease and phosphatase inhibitors (5 µg/ml Pepstatin A, 5 µΜ Leupeptin, 1mM AEBSF, 1 µg/ml microcystin) as previously described (Louneva et al., 2008; Pensalfini et al., 2020). Samples were adjusted to contain 1.25 M sucrose and sequentially overlaid with 1.0 M sucrose solution containing 0.1 mM CaCl_2_, followed by a layer of homogenization buffer. Tubes were centrifugated at 100,000 x g for 3 hours at 4°C using a SW55Ti rotor (Beckman Coulter). Synaptosomes enriched in the interface between 1.0-1.25 M sucrose were collected and split equally between synaptic vesicle endocytosis analyses (see below) and biochemical analyses. For biochemistry, the synaptosomes were washed twice with ice-cold 0.1 mM CaCl_2_, followed by centrifugation at 50,000 x g for 30 minutes at 4°C in a SW55Ti rotor. Washed pellets were dissolved in 8M Urea prior to further biochemical analyses.

### 2.8. Synaptic vesicle endocytosis

Synaptosomes from non-Tg and Thy1-APPL1 mouse hippocampi (n=4-6 mouse each genotype) were used for synaptic vesicle endocytosis analysis (Daniel et al., 2012; Sellgren et al., 2019; Shekarabi et al., 2019). Synaptosomes from the 1.0-1.25 M sucrose interface were resuspended in sucrose/EDTA/Tris buffer (SET buffer: 0.32 M sucrose, 1 mM EDTA, 5 mM Tris (pH 7.4)) and centrifugated at 20,000 x g for 10 minutes at 4°C in a SW55Ti rotor, followed by resuspension in SET buffer. The resuspended synaptosomes were aliquoted and slowly frozen with 5% DMSO in SET buffer to preserve integrity as described (Daniel et al., 2012). On the day of the assay, aliquots of synaptosomes were thawed in a 37°C water bath for 80 seconds and mixed with 1 ml of ice-cold SET buffer, followed by centrifugation at 18,900 x g for 2 minutes at 4°C to pellet the synaptosomes. The synaptosome pellets were again suspended in 300 µl of SET buffer with a wide-orifice 1 ml pipette tip. After protein quantification, 2 µg of synaptosomes were suspended in SET buffer containing 250 µM dithiothreitol to prevent clumping and loaded per well into a polyethyleneimine-coated 96-well glass-bottom microplate. The 96-well plate with synaptosomes was then centrifugated in a microplate centrifuge (Sigma, 4-15C) at 1,500 x g for 30 minutes at 4°C to seed the synaptosomes onto the plate before incubation in HBK buffer (HEPES-buffered Krebs-like buffer; 143 mM NaCl, 4.7 mM KCl, 1.3 mM MgSO_4_, 1.2 mM CaCl_2_, 20 mM HEPES, 0.1 mM NaH_2_PO_4_ and 10 mM D-glucose, pH 7.4) at 37°C for 15 minutes. The synaptosomes were then labeled with 5 µM of CellTracker^TM^ Green CMFDA (5-chloromethyl fluorescein diacetate; Thermo Fisher; Cat# C7025) to label viable synaptosomes for 30 minutes at 37°C, followed by an 8-minute pulse with 4 µM of the endocytic dye FM^TM^4-64 (*N*-(3-Triethylammoniumpropyl)-4-(6-(4-(Diethylamino) Phenyl) Hexatrienyl) Pyridinium Dibromide, Thermo Fisher; T13320). Labeled synaptosomes were then washed with 1 mM of Advasep-7 (Biotium, 70029) for 2 minutes to remove extracellular dye, before fixation with 2% PFA in HBK buffer for 30 minutes on ice, followed by imaging at 40X magnification (with 3X zoom) using a LSM880 confocal microscope. For quantification, 25-30 synaptosome images were taken from each mouse, and the intensity of FM^TM^4-64 and CMFDA within synaptosomes was measured with Cell Profiler (https://cellprofiler.org) to obtain the ratio of FM^TM^4-64/CMFDA per synaptosome in non-Tg and Thy1-APPL1 mice (n=4-6). Additional immunofluorescence staining with anti-rab5 antibody (Abcam, 1:200) was performed after CMFDA intake while avoiding dislodging the attached synaptosomes. The intensity of the rab5 and CMFDA signals was measured with ImageJ to determine the ratio of rab5/CMFDA in non-Tg and Thy1-APPL1 mice (n=3).

### 2.9. GTP-agarose beads pull down assay

Synaptosomes isolated as described above from non-Tg and Thy1-APPL1 mouse hippocampus (n=3 mice per genotype) were used for a GTP-agarose beads pull down assay (Zhang et al., 2013; Fang et al., 2017; Pensalfini et al., 2020). Briefly, synaptosome pellets were resuspended in GTP-agarose lysis/wash buffer (50 mM Tris-HCl, pH7.5; 250 mM NaCl, 5mM Mg Acetate, 0.5% Triton X-100, and protease inhibitors), incubated on ice for 30 minutes, then sonicated prior to a BCA protein assay. 200 µg of the synaptosome lysates were added into 300 µl prewashed GTP-agarose beads (Sigma, cat# G9768). The synaptosome and bead mixtures were incubated overnight at 4°C with rotation, then washed three times with wash buffer before being boiled for 10 minutes in 2x Laemmli sample buffer. As reference, 10 µg of synaptosome lysate was used as “Input” for western blot analysis as described above.

### 2.10. Novel Object Recognition

Novel object recognition was performed according to our previous publications (Pensalfini et al., 2020; Jiang et al., 2022). A multiple unit open field with four activity chambers and a digital video camera were used to record the activity of the mice inside each chamber. 12-13-month-old non-Tg and Thy1-APPL1 mice (n=10 for both genotypes) were used for the test. Mice were habituated by freely exploring the chamber for 5 minutes per day for two days. On day three, a training session was conducted by placing an individual mouse together with two identical objects (familiar) for 10 minutes, followed by one familiar and one novel object for 10 minutes after a three-hour interval. The recorded videos were analyzed with CowLog 3 (Hänninen and Pastell, 2009) to determine the exploration time of each mouse. The results were presented as Recognition Index (RI), defined as time exploring the novel object versus the sum of the time exploring both the novel and familiar objects, as previous described (Pensalfini et al., 2020; Jiang et al., 2022).

### 2.11. Experimental Design and Statistical Analysis

All quantitative data were subjected to a two-tailed unpaired Student’s t-test for a single comparison and a one-way ANOVA analysis for multiple comparisons, with post-hoc Tukey’s analysis, using GraphPad Prism 8.0.1. Data is represented in bar graphs as the Mean ± SEM, with individual data points for each mouse in the study group. Statistical significance is represented by asterisks: *p<0.05, **p<0.01, ***p<0.001. Sample sizes were determined based on similar experimental procedures in our previous studies (Pensalfini et al., 2020; Jiang et al., 2022). Initial age-studies showed that the basal forebrain cholinergic neurons (BFCN) loss in the Thy1-APPL1 mice was first apparent at 12-13 months of age; therefore, unless specifically stated in the text, most presented findings are of non-Tg and Thy1-APPL1 mice at 12-13 months of age.

## 3. Results

### 3.1. APPL1 overexpression in the brains of transgenic mice

APPL1 transgenic lines under the Thy1 promoter were generated, and an experimental line was chosen and backcrossed prior to expansion and experimentation (Thy1-APPL1). APPL1 overexpression was confirmed by western blot analysis (Fig. 1A). Greater APPL1 immunolabeling was seen broadly in multiple brain regions (including cortex, hippocampus, cerebellum) when comparing Thy1-APPL1 with non-Tg mice (Fig. 1A-B), with a neuronal expression shown by coincident immunolabeling of APPL1 (green) and NeuN (red) (Fig.1C; cortex and hippocampus). Increased immunolabeling for APPL1 is evident in neurons when comparing Thy1-APPL1 and non-Tg brain tissue (Fig. 1C, comparing yellow APPL1 labeling in NeuN-positive neurons, yellow arrows). Non-neuronal cells identified by DAPI nucleus staining showed similar levels of APPL1 immunolabeling in both Thy1-APPL1 and non-Tg brain tissue (Fig. 1C, white arrow heads). Western blot analysis of brain cortex homogenate across three age cohorts ranging from 4-5 to 12-13 months of age showed an approximate doubling of brain APPL1 levels in the Thy1-APPL1 mice compared to non-Tg littermates (Fig. 1D-E; n=3-5 for non-Tg and Thy1-APPL1; *t(6)=4.550, p=0.00392* for 4-5 month-old mice; *t(6)=11.5, p=0.000025* for 7-9 month-old mice; *t(6)=6.387, p=0.000693* for 12-13 month-old mice; Two-tailed, unpaired t-test). Detected with a pan-rab5 antibody that does not differentiate active from inactive forms of the protein, rab5 protein levels in brain homogenates were unchanged when comparing APP1 overexpressing mice to non-Tg (Fig. 1D-E). Thus, Thy1-APPL1 mice show neuronal specific APPL1 overexpression across many brain regions, an increase that does not result in a change in overall brain rab5 protein levels.

**Figure 1.**
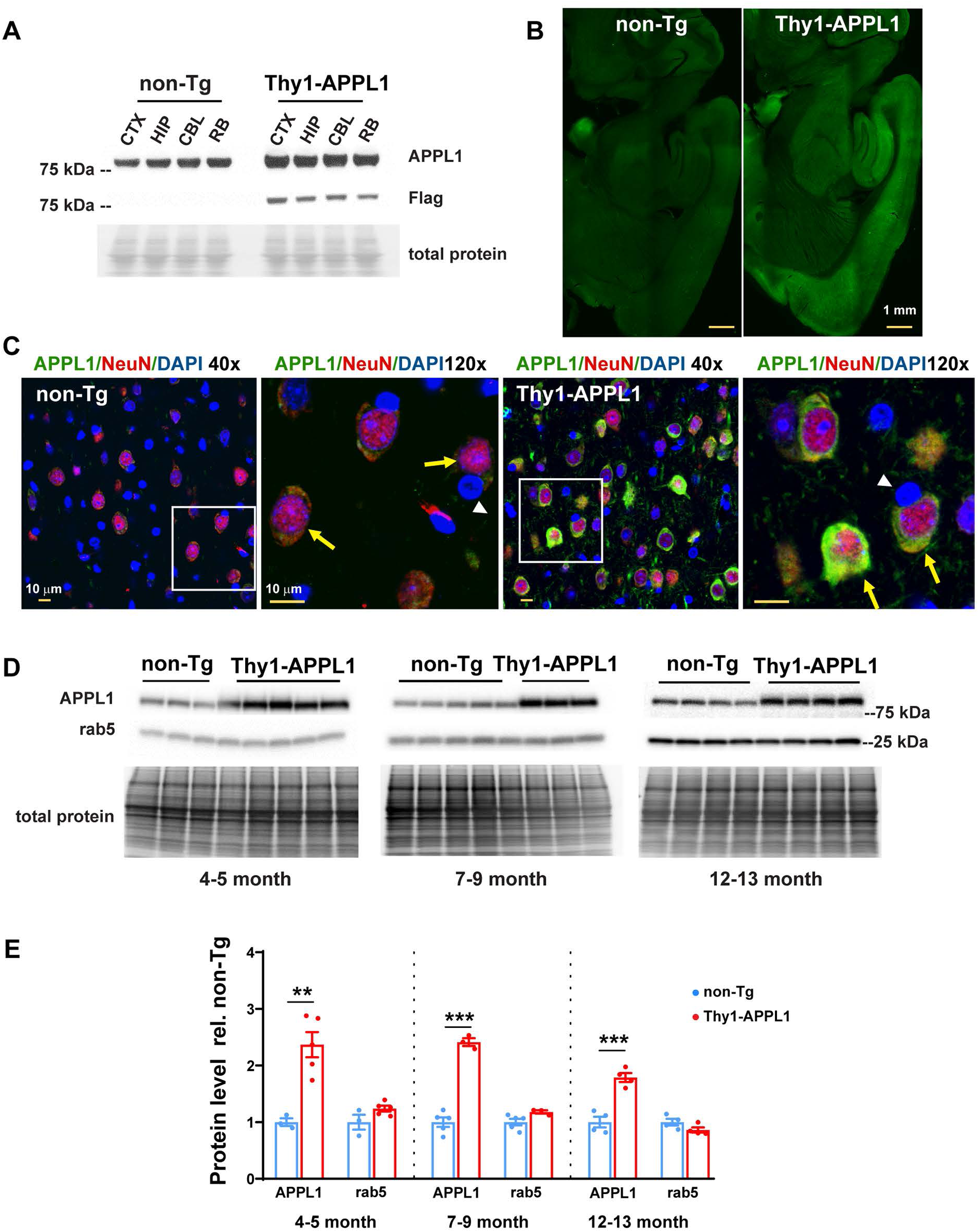
APPL1 transgene expression in the brain. ***A.*** Representative western blot comparing APPL1 expression in various brain regions (CTX-cortex; HIP-hippocampus; CBL-cerebellum; RB-remaining brain) from 4–5-month-old non-transgenic littermate (non-Tg) and Thy1-APPL1 transgenic (Thy1-APPL1) mice. Over-expressed flag-tagged APPL1-proteins were detected in all brain regions in Thy1-APPL1 mice. ***B*.** Representative brain section immunolabeling with an anti-APPL1 antibody at low magnification (scale bar 1mm) demonstrates the APPL1 over-expression patterns in various brain regions of Thy1-APPL1 mice. ***C.*** Representative immunofluorescent images at low (40x) and high (120x) magnification show APPL1 (green) and NeuN (red) with DAPI (blue) staining in cortex layer V (scale bar, 10 µm; yellow arrows indicate neurons, white arrowheads identify non-neuronal cells; genotype as indicated) ***D*.** Western blots probed with anti-APPL1 and anti-rab5 antibodies of both non-Tg and Thy1-APPL1 brain homogenate prepared from mice of the indicated ages. ***E.*** Western blot analysis quantification showing increased APPL1 in the brain homogenates of Thy1-APPL1 mice compared to non-Tg mice at the indicated ages. \*\**p<0.01,* \*\*\**p<0.001,* Two-tailed, unpaired t-test. Data shown as Mean ± SEM.

### 3.2. Thy1-APPL1 transgenic mice have early endosomal abnormalities

Our previous study showed that APPL1 is a molecular intermediary between APP-βCTF-driven endosomal alterations and rab5, required for increased APP-βCTF levels to drive such early endosome dysfunctions as enlargement and disrupted trafficking in a cell model (Kim et al., 2016). One goal of the current study using the Thy1-APPL1 mice was to determine whether an increase in APPL1 is sufficient to drive the same range of early endosome pathologies *in vivo*, directly testing whether APPL1 can play a primary role in AD-relevant early endosomal alterations. Immunolabeling with an anti-rab5 antibody that only detects the active, early endosome-membrane associated, GTP-bound state of rab5 (rab5-GTP) (Mani et al., 2016; Pensalfini et al., 2020) was done comparing cortical pyramidal neurons between Thy1-APPL1 and non-Tg mice (Fig. 2A). A significantly higher level of active rab5 immunolabeling was seen in cortical layer V neurons of Thy1-APPL1 mice compared to non-Tg littermates. Representative immunolabeling at low magnification of 12-month-old non-Tg and Thy1-APPL1 mice shows the coincidence of rab5-GTP (red) and APPL1 (green) (Fig. 2A, left hand panels). Higher magnification of a single neuron illustrates typical rab5-GTP and APPL1 immunolabeling in both genotypes (Fig. 2A). There was also significantly higher colocalization of APPL1 with rab5-GTP in Thy1-APPL1at 12-13 month of age, with *Pearson’s R=0.161* in non-Tg vs *R=0.303* in Thy1-APPL1 mice (n=4 each for non-Tg and Thy1-APPL1 mice, *t(6)=3.986, p=0.0072*, Two-tailed, unpaired t-test), consistent with overexpressed APPL1 leading to the recruitment of rab5 to early endosomes.

**Figure 2.**
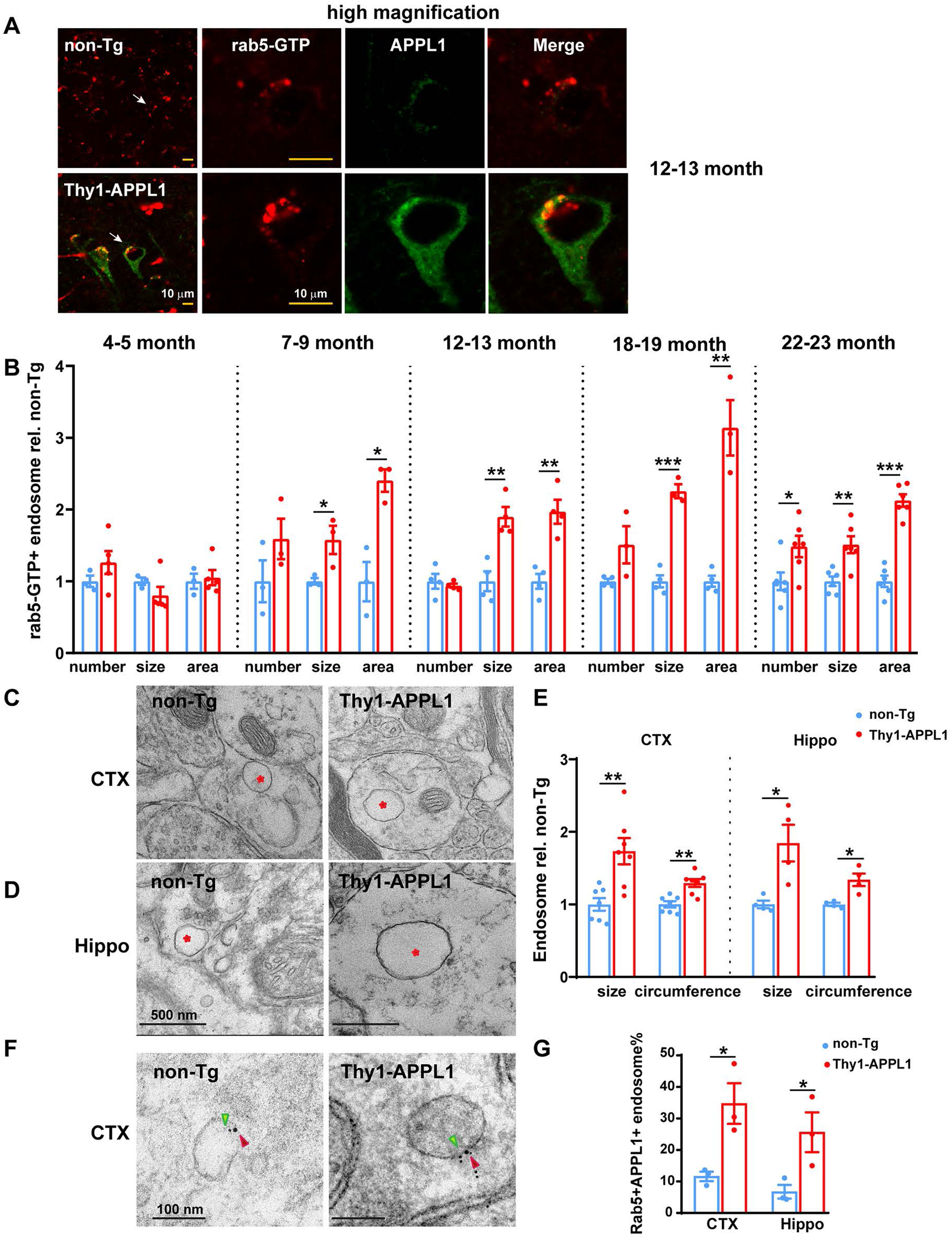
Early endosome alterations in Thy1-APPL1 mice. ***A.*** Representative immunofluorescent images of active, GTP-bound rab5 (rab5-GTP; red) and APPL1 (green) in neurons (layer V, pre-frontal cortex) of non-Tg and Thy1-APPL1 mice at 12-13 months of age. Shown are images at lower magnification (40X, merged). The white arrow indicating the neuron shown at higher magnification for individual and merged immunofluorescent signal. Scale bar 10 µm. ***B*.** Quantification of the average number, size, and total area of Rab5-GTP immunolabeled endosomes per cortical neuron in non-Tg versus Thy1-APPL1 mice at the indicated ages. Representative electron microscopy images containing dendritic profiles in layer V of pre-frontal cortex (***C***) and hippocampus (***D***) regions, and (***E***) quantification of average size and circumference of endosomes in both cortex and hippocampi showing the enlargement in Thy1-APPL1 compared to non-Tg mice aged at 12-13 months (red asterisk indicating endosome; 7 mice each genotype for cortex and 4 mice each genotype for hippocampi were used for quantification. Scale bar, 500 nm). Sixty images/mouse were quantified in (***E***); for cortex sections, a total of 2635 endosomes from 14 mice were counted, averaging 188 endosomes per mouse; For hippocampal sections, a total of 1594 endosomes were counted from 8 mice, with an average of 199 endosomes per mouse. ***F.*** Representative immuno-electron microscopy images containing dendritic profiles in layer V of pre-frontal cortex labeled with rabbit anti-rab5 and mouse anti-APPL1 followed by gold-conjugated secondary anti-rabbit (6 nm, green arrow) and anti-mouse (10 nm, red arrow) in non-Tg and Thy1-APPL1 mice. Scale bar 100 nm. ***G.*** Percentage of rab5/APPL1 double-positive endosomes versus total rab5-positive endosomes in the dendritic region of cortex and hippocampus significantly increased in Thy1-APPL1 mice (3 mice per genotype. 35-40 images/mouse were used for counting; for cortex sections, a total of 130 endosomes were counted from 6 mice, with an average of 22 endosomes per mouse; for hippocampal sections, a total of 121 endosomes were counted from 6 mice, with an average of 20 endosomes per mouse). \**p<0.05,* \*\**p<0.01,* \*\*\**p<0.001,* Two-tailed, unpaired t-test. Data is shown as Mean ± SEM.

The quantitative data across the life-span of non-Tg and Thy1-APPL1 mice are summarized in Fig. 2B. (n=3-6 each age and genotype; *t(6)=1.198, p=0.276; t(6)=1.201, p=0.273; t(6)=0.318, p=0.761;* for number, size and area of 4-5 month-old mice; *t(4)=1.457, p=0.219; t(4)=2.844, p=0.0467; t(4)=4.467, p=0.0111* for number, size and area of 7-9 month-old mice, respectively; *t(6)=0.577, p=0.585; t(6)=4.614, p=0.00364; t(4)=4.930, p=0.00263* for number, size and area of 12-13 month-old mice; *t(5)=2.307, p=0.0691; t(5)=9.717, p=0.000196; t(5)=6.356, p=0.00142;* for number, size and area of 18-19 month-old mice; *t(10)=2.515, p=0.0306; t(10)=3.737, p=0.00386; t(10)=9.818, p=0.000002;* for number, size and area of 22-23 month-old mice; Two-tailed, unpaired t-test). Interestingly, the youngest age group (4–5-month-old) Thy1-APPL1 mice did not show an increase in rab5-GTP early endosome number or size compared to non-Tg, although APPL1 is overexpressed at this age (Fig. 1D-E). While early endosome size and, correspondingly, early endosome area/cell were increased in all of the older age groups, the number of early endosomes was not found to be increased until the oldest cohort of Thy1-APPL1 mice (Fig.2B, age 22-23 months). Age-dependent endosome alterations were also demonstrated in APOE4 mice (Nuriel et al., 2017) and Down Syndrome’s mouse models (Jiang et al., 2016), suggesting that endosome homeostasis may be maintained at younger ages despite a perturbation that ultimately drives endosomal dysfunction as the brain ages.

The endosomal enlargement due to APPL1 overexpression was confirmed by electron microscopy (EM) in 12-13 month-old Thy1-APPL1 compared to non-Tg mice, where both endosomal size and circumference were significantly increased in layer V cells of prefrontal cortex (Fig. 2C, 2E; n=7 for non-Tg and Thy1-APPL1 mice, *t(12)=3.64, p=0.0034*; *t(12)=4.239, p=0.0012* for size and circumference, respectively; Two-tailed, unpaired t-test) and the CA1 region of the hippocampus (Fig. 2D-E; n=4 for non-Tg and Thy1-APPL1 mice, *t(6)=3.327, p=0.0173*; *t(6)=3.83, p=0.00866* for size and circumference, respectively; Two-tailed, unpaired t-test). Increased association of APPL1 with rab5-positive endosomes has been previously shown in cortical neurons from both AD and DS human brain (Kim et al., 2016), consistent with the increased co-localization of APPL1 and rab5-GTP seen in the Thy1-APPL1 mice (Fig. 2A). To further substantiate this co-localization at the early endosome, we performed immuno-EM of cortical and CA1 hippocampal brain sections of 12-13-month-old mice using a rabbit anti-rab5 and a murine anti-APPL1 antibody, followed by species-specific gold-conjugated secondary antibodies (anti-rabbit, 6 nm, green colored; anti-mouse, 10 nm, red colored). EM images from fields containing dendritic profiles were collected and the number of gold-particle-positive endosomal vesicles was manually counted. The ratio of rab5/APPL1 double-positive endosomes vs. total rab5 positive endosomes was significantly higher in Thy1-APPL1 compared to non-Tg mice in both brain regions (Fig. 2F-G; n=3 for non-Tg and Thy1-APPL1 mice, *t(4)=3.492, p=0.0251*; *t(4)=2.825, p=0.0476* for cortex and CA1, respectively; Two-tailed, unpaired t-test).

### 3.3. Endocytic abnormalities in hippocampal synaptosomes of Thy1-APPL1 mice

Given that our previous studies demonstrated that rab5 hyper-activation leads to synaptic dysfunction and deficits in memory and learning (Kaur et al., 2014; Jiang et al., 2016; Pensalfini et al., 2020) and that endocytic activity plays a critical role in synaptic function (Overhoff et al., 2021), we sought to confirm that the overexpression of APPL1 in the Thy1-APPL1 mouse leads to differences in endocytic function in isolated synaptosomes. Independent of genotype, synapse enrichment was confirmed by the higher level of synaptophysin in the synaptosome preparations compared to hippocampal homogenates (Fig. 3A and additional data not shown; n=6 per genotype, at the age of 12-13 months). Consistent with the neuronal expression of APPL1 in the transgenic mice (Fig. 1C), the increase of APPL1 protein in the synaptosomes prepared from Thy1-APPL1 mice was found to be greater than that seen in overall brain homogenate (Fig. 3A-B, 3.9-times non-Tg in synaptosomes compared to 1.9-times in homogenate, n=6, *t(10)=5.36, p=0.000319,* Two-tailed, unpaired t-test). Total rab5 protein levels in both homogenates (also in Fig. 1D-E) and synaptosomes were unaffected by APPL1 overexpression (Fig. 3A-B). While APP-βCTF levels were similar in Thy1-APPL1 and non-Tg mouse brain homogenate, APPL1 overexpression led to an increase in APP-βCTF levels within synaptosomes (Fig. 3A-B; n=6 per genotype; *t(10)=3.054, p=0.0122*; Two-tailed, unpaired t-test). Given that the APP-βCTF is the immediate precursor of Aβ, we determined brain Aβ levels using a sandwich ELISA that detects the endogenous, murine Aβ (Morales-Corraliza et al., 2009; Jiang et al., 2016) in 7-9- and 12–13-month-old mice. The levels of brain Aβ40 and Aβ42 were found to be unchanged when comparing Thy1-APPL1 to non-Tg mice brain homogenate (data not shown).

**Figure 3.**
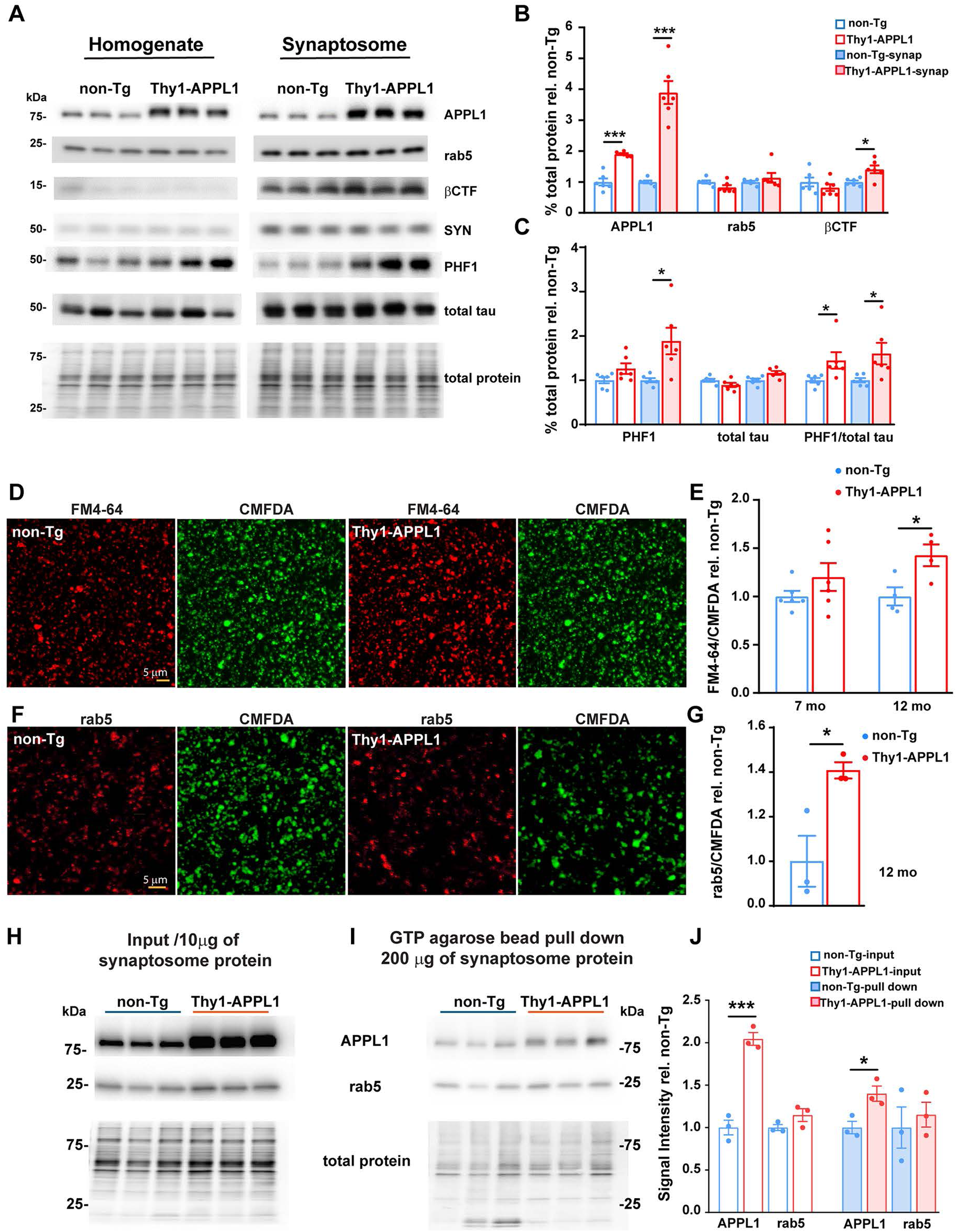
Western blot analysis and synaptic vesicle endocytosis (SVE) revealed the abnormalities in hippocampal synaptosomes of Thy1-APPL1 mice. ***A.*** Representative western blots from one of the two experiments showing various protein markers in hippocampal homogenates and hippocampal synaptosome preparations from non-Tg and Thy1-APPL1 mice (n=6 each genotype). Quantitation of these western blots (***B-C***) shows significantly higher levels of APPL1, βCTF, PHF1 and the ratio of PHF1/total tau in synaptosomes of Thy1-APPL1 compared to non-Tg mice at 12–13 months of age. ***D.*** Representative fluorescent images of internalized FM4-64 (red) with corresponding CMFDA labeling of total synaptosomes (green) using hippocampal synaptosomes prepared from non-Tg and Thy1-APPL1 mice at 12-13 months of age. (***E***) The ratio of internalized FM4-64 to CMDFA was significantly increased in hippocampal synaptosomes of Thy1-APPL1 mice at 12-13 months, but not in 7 months of age, indicating elevated endocytosis in older Thy1-APPL1 mice compared to non-Tg. ***F.*** Representative images of CMFDA (green) labeling followed by anti-rab5 (red) immuno-labeling and (***G***) quantification of the intensity of rab5 to CMFDA showing an increase in rab5 immunosignal per CMFDA labeled synaptosome in hippocampal synaptosomes of Thy1-APPL1 mice at 12-13 months of age. Scale bar, 5 µm, \**p<0.05,* \*\*\**p<0.001,* Two-tailed, unpaired t-test. Data shown as Mean ± SEM. Western blots of hippocampal synaptosome (10 µg of synaptosome protein, input) (***H***), and GTP-agarose pull down of hippocampal synaptosome (200 µg of synaptosome protein) from non-Tg and Thy1-APPL1 mice (***I***) probed with anti-APPL1 and anti-rab5 antibodies, total protein stained by Revert^TM^ 700 Total protein Stain are also shown at the bottom of the (***H***) and (***I***). The APPL1 and rab5 band density in both input and pull down against total protein in relation with non-Tg are presented in ***(J)***, \**p<0.05,* \*\*\**p<0.001,* one-way ANOVA, Data is shown as Mean ± SEM.

Phosphorylated tau (identified with PHF1) (Pensalfini et al., 2020) was also increased in hippocampal synaptosomes of Thy1-APPL1 mice compared to non-Tg controls (Fig. 3A and 3C; n=6 for per genotype; *t(10)=2.922, p=0.0152* for synaptosome PHF1; *t(10)=2.309, p=0.0436* for PHF1/total tau in brain homogenate; *t(10)=2.432, p=0.0353* for PHF1/total tau in synaptosome homogenate; Two-tailed, unpaired t-test), an increase in PHF1-reactivity comparable to the increase seen following neuronal rab5 overexpression (Pensalfini et al., 2020). These findings in Thy1-APPL1 mouse further argue that endosomal pathway disruption can contribute to tau pathobiology in AD (Knopman et al., 2021).

To directly investigate whether endocytosis was altered by the increase in APPL1 expression in the Thy1-APPL1 mice, total synaptosomes were identified with the fluorescent dye CMFDA (Daniel et al., 2012; Sellgren et al., 2019). *Ex vivo* synaptosome endocytic uptake was determined using a second fluorescent marker, FM4-64, following its internalization (Daniel et al., 2012; Sellgren et al., 2019). Confocal images of both FM4-64 and CMFDA were collected and the ratio of FM4-64 to CMFDA in synaptosome was quantified. A significantly greater ratio of FM4-64/CMFDA was observed in hippocampal synaptosomes isolated from Thy1-APPL1 compared to non-Tg mice at 12 months of age (Fig. 3D-E, n=4 for non-Tg and Thy1-APPL1 mice; *t(6)=2.886, p=0.0278;* Two-tailed, unpaired t-test), while the ratios of FM4-64/CMFDA remain unchanged between non-Tg and Thy1-APPL1 mice at the age of 7 months (Fig. 3E, n=6; *t(10)=1.294, p=0.225;* Two-tailed, unpaired t-test). This age-dependent increase in synaptosomal endocytic uptake is consistent with the age-dependent increase in neuronal early endosome size seen in the Thy1-APPL1 mice (Fig. 2B). As determined by immunocytochemistry using an anti-rab5 antibody (Abcam, 1;200), a significantly higher ratio of rab5/CMDFA was also found in Thy1-APPL1 mice at the age of 12 months (Fig. 5F-G, n=3 for non-Tg and Thy1-hAPPL1 mice; *t(4)=3.403, p=0.0272;* Two-tailed, unpaired t-test), supporting the idea that rab5 endosomal recruitment in the Thy1-APPL1 mouse leads to greater endocytosis. These findings support our hypothesis that APPL1 acts as a molecular interface connecting rab5 and APP-βCTFs at the early endosome.

We and others have previously used GTP-agarose affinity columns to isolate rab5 from brain tissue and cells (Xu et al., 2016; Fang et al., 2017; Pensalfini et al., 2020). In this study, we used GTP-agarose-affinity to isolate rab proteins and to co-isolate rab5-associated APPL1 (Miaczynska et al., 2004; Mao et al., 2006; Zhu et al., 2007; Hiragi et al., 2022) to confirm within neurons an increased association of APPL1 and rab5 in the Thy1-APPL1 mouse compared to non-Tg. Consistent with the increased level seen in brain homogenate (Fig. 1), APPL1 was more abundant in the synaptosome preparation from Thy1-APPL1 mice compared to non-Tg (Fig. 3H, J). Following GTP-agarose isolation of rab proteins, APPL1 was co-isolated from both Thy1-APPL1 and non-Tg mice. However, more APPL1 in the GTP-agarose pull-down was seen in the Thy1-APPL1 synaptosomes than in those prepared from non-Tg mice (Fig. 3I and J) (n=3 for non-Tg and Thy1-APPL1 mice; for APPL1 Input, t*(8)=9.081, p<0.0001;* for APPL1 pull down, *(8)=3.472, p=0.0168;* One-way ANOVA). Similar levels of rab5 were isolated from both genotypes using the GTP-agarose pull-down, consistent with our findings showing that total rab5 protein levels are not increased in the Thy1-APPL1 mouse, both in brain homogenates and isolated synaptosomes (Fig. 3A-B, H and J). The greater amount of APPL1 isolated from Thy1-APPL1 mouse synaptosomes using the rab GTP-affinity column is consistent with previous findings of APPL1 and rab5 directly interacting (Miaczynska et al., 2004; Mao et al., 2006; Zhu et al., 2007; Hiragi et al., 2022), and supports our interpretation that the synaptosomal endocytic differences seen in Thy1-APPL1 result from APPL1 over-expression leading to enhanced rab5 activation due to an enhanced interaction between the two molecules.

### 3.4. Loss of BFCNs and endosome accumulation within neurons in Thy1-APPL1 mice

Neuronal endosomal dysfunction drives the loss of cholinergic neurons in the MSN due to a loss of trophic support mediated by signaling endosomes (Salehi et al., 2003; Howe and Mobley, 2004; Jiang et al., 2016; Xu et al., 2018; Chen and Mobley, 2019; Pensalfini et al., 2020; Jiang et al., 2022). Consistent with the endosomal alterations seen in the Thy1-APPL1 model, the number of ChAT-immunoreactive neurons in the MSN of Thy1-APPL1 mice showed an age-dependent loss compared to non-Tg mice (Fig. 4A-B, n=42 for total of non-Tg mice and n=41 for total of Thy1-APPL1 mice; *F(1, 55)=6.01, p=0.017,* Linear regression), with reduced numbers of cholinergic neurons first detected at 12 months of age (Fig. 4C, *t(25)=2.316, p=0.0291* for 12-13 month old mice; *t(7)=2.399, p=0.0475* for 18 month old mice; *t(9)=3.482, p=0.00692* for mice over 23 month old; Two-tailed, unpaired t-test).

**Figure 4.**
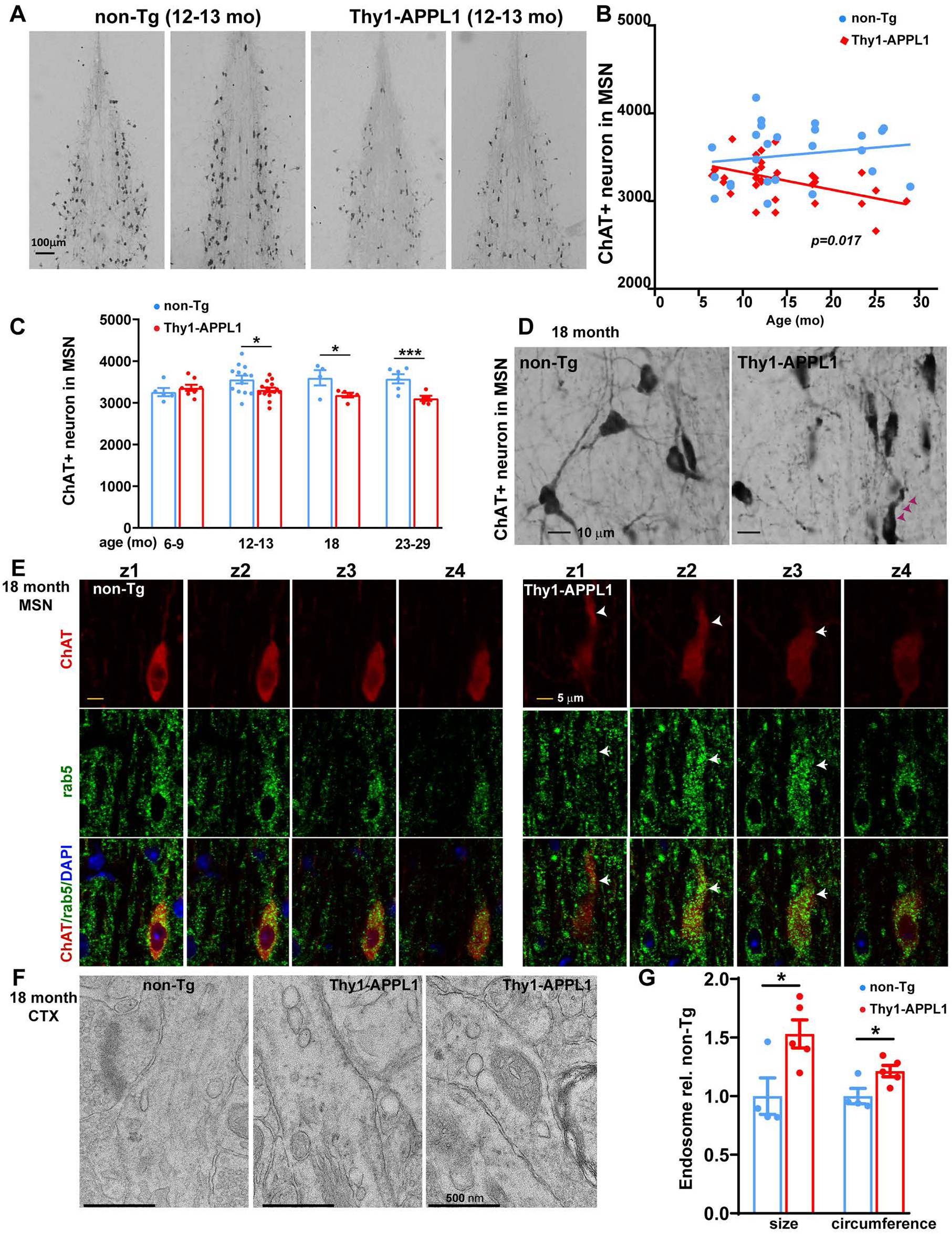
Loss of cholinergic neurons in the medial septal nucleus (MSN) of Thy1-APPL1 mice and accumulation of endosomes in Thy1-APPL1 mice. ***A.*** Representative light microscopy images comparing ChAT immunoreactive neurons in the MSN of non-Tg and Thy1-APPL1 mice at 12-13 months of age. Scale bar, 100 µm. ***B.*** Plots of number of ChAT^+^ neurons counted stereologically versus age of mice showing significant difference between the slopes of non-Tg and Thy1-APPL1 mice (*F(1, 55)=6.01, p=0.017,* Linear regression). ***C.*** Number of ChAT -immunoreactive neurons in the MSN of non-Tg and Thy1-APPL1 mice at the indicated ages (n = 42 for non-Tg and 41 for Thy1-APPL1 mice; quantitative stereology as described in Methods). ***D.*** Representative images of ChAT-immunoreactive neurons at higher magnification in 18-month-old non-Tg and Thy1-APPL1 mice (arrow heads indicate dystrophic neurites). Scale bar, 5 µm. ***E***. Representative z-stacked series fluorescence images of ChAT-(red) and rab5-immunoreactive (green) neuron in the MSN of a non-Tg and Thy1-APPL1 mice at 18 months of age, (arrow heads indicating a dystrophic neurite). Scale bar, 5 µm. \**p<0.05,* \*\*\**p<0.001,* Two-tailed, unpaired t-test. Data presented as Mean ± SEM. ***F.*** Representative EM images of 18-month-old mice showing an accumulation of endosomes in cortical dendrites of Thy1-APPL1 mice compared to non-Tg mice (scale bar, 500 nm). ***G.*** Quantification of average size and circumference of endosomes in the cortical dendritic area of non-Tg and Thy1-APPL1 mice at the age of 18 months (n=4 for non-Tg, and n=5 for Thy1-APPL1 mice; 60 cortical section images/mouse, a total of 982 endosomes were counted in Thy1-APPL1mice, with an average of 196 endosomes/mouse counted. 630 total endosomes were counted from non-Tg mice, with averaging 157 endosomes/mouse,), \**p<0.05,* Two-tailed, unpaired t-test. Data shown as Mean ± SEM.

Abnormal, tortuous proximal dendrites and shrunken cholinergic cell soma (Fig. 4D, arrowheads) were also seen in the MSN neurons of aged Thy1-APPL1 compared to non-Tg mice (18 month), as has been reported in multiple AD-like models with endosomal alterations (Cataldo et al., 2000; Cataldo et al., 2003; Salehi et al., 2006; Choi et al., 2009; Jiang et al., 2016; Pensalfini et al., 2020; Jiang et al., 2022). Fig. 4E shows representative Z-stacked images of immunolabeling rab5-positive early endosomes within cholinergic neurons in a Thy1-APPL1 compared to a non-Tg mouse, both at 18-months of age. Accumulation of early endosomes (arrowheads) was apparent in the ChAT-immunoreactive cholinergic neurons of the Thy1-APPL1 relative to non-Tg mice. Representative EM images of 18-month-old mice also demonstrated an accumulation of endosomes (Fig. 4F) as well as a significant endosome size increase in cortical dendritic areas of Thy1-APPL1 mice compared to non-Tg (Fig. 4G; n=4 for non-Tg, and n=5 for Thy1-APPL1 mice; *t(7)=2.753, p=0.0284* for the size; *t(7)=2.701, p=0.0306* for the circumference). This EM finding is consistent with the previous immunolabeling documenting an increase in active rab5 early endosomes in neurons of the Thy1-APPL1 mice (see Fig. 2).

### 3.6. Hippocampal synaptic plasticity impairments in Thy1-APPL1 mice

Alteration of hippocampal synaptic plasticity has been shown in AD-related mouse models (Marchetti and Marie, 2011; Dietrich et al., 2018), and APPL1 has been shown to be a regulator of hippocampal synaptic plasticity (Fernandez-Monreal et al., 2016; Wang et al., 2016; Formolo et al., 2022; Hua et al., 2023). We have also previously demonstrated that over-activation of rab5 resulting from neuronal rab5 overexpression leads to hippocampal-dependent memory deficits (Pensalfini et al., 2020). Long-lasting changes in synaptic plasticity measured as LTP and LTD are key mechanisms upon which learning and long-term memory are built (Avchalumov and Mandyam, 2021). Greater rab5 activation has been linked with both LTP deficit (Pensalfini et al., 2020) and LTD impairments (Hausser and Schlett, 2019; Pensalfini et al., 2020). To determine whether APPL1 overexpression in the Thy1-APPL1 mouse altered hippocampal plasticity, we measured LTP and LTD of CA1hippocampal slices of Thy1-APPL1 and non-Tg mice (Fig. 5; n=5 for both genotypes at ages 7-9 and 12-13 months). In the younger age cohort (7-9 months), Thy1-APPL1 mice did not show differences compared to non-Tg in LTP following TBS stimulation (Fig. 5A-C). However, we found a significant LTP impairment in 12-13 month-old Thy1-APPL1 mice (Fig. 5E: (*F(1, 236)=6.696, p=0.0103* for the slope, Linear regression) and Fig. 5F: (*t(8)=2.565, p=0.0334* for 1 minute time point*; t(8)=3.237, p=0.0237* for 40 minutes time point; *t(8)=3.722, p=0.0175* for 80 minutes time point; Two-tailed, unpaired multiple t-test), without apparent difference in their basal neurotransmission between the genotypes (Fig. 5D). Furthermore, post-synaptic potential (fEPSP) slopes indicated suppressed LTD upon induction by low frequency stimulation (1 Hertz for 900 seconds) in 12-13 month-old Thy1-APPL1 mice compared to their age-matched non-Tg littermates (Fig. 5H: (*F(1, 127)=14.83, p=0.0002* for the intercepts, Linear regression and Fig. 5I: (*t(8)=5.277, p=0.000749* for 50 minutes time point; Two-tailed, unpaired t-test). Thus, we show both loss of BFCNs as well as disruption of hippocampal CA1 LTP and LTD as the Thy1-APPL1 mice age. Consistent with these electrophysiological findings, hippocampal-dependent memory assessed by Novel Object Recognition (NOR) showed impairment in Thy1-APPL1 mice at 12-13 months of age (Fig. 5J; n=10 for non-Tg and Thy1-APPL1 mice; *t(18)=2.187, p=0.042*; Two-tailed, unpaired t-test).

**Figure 5.**
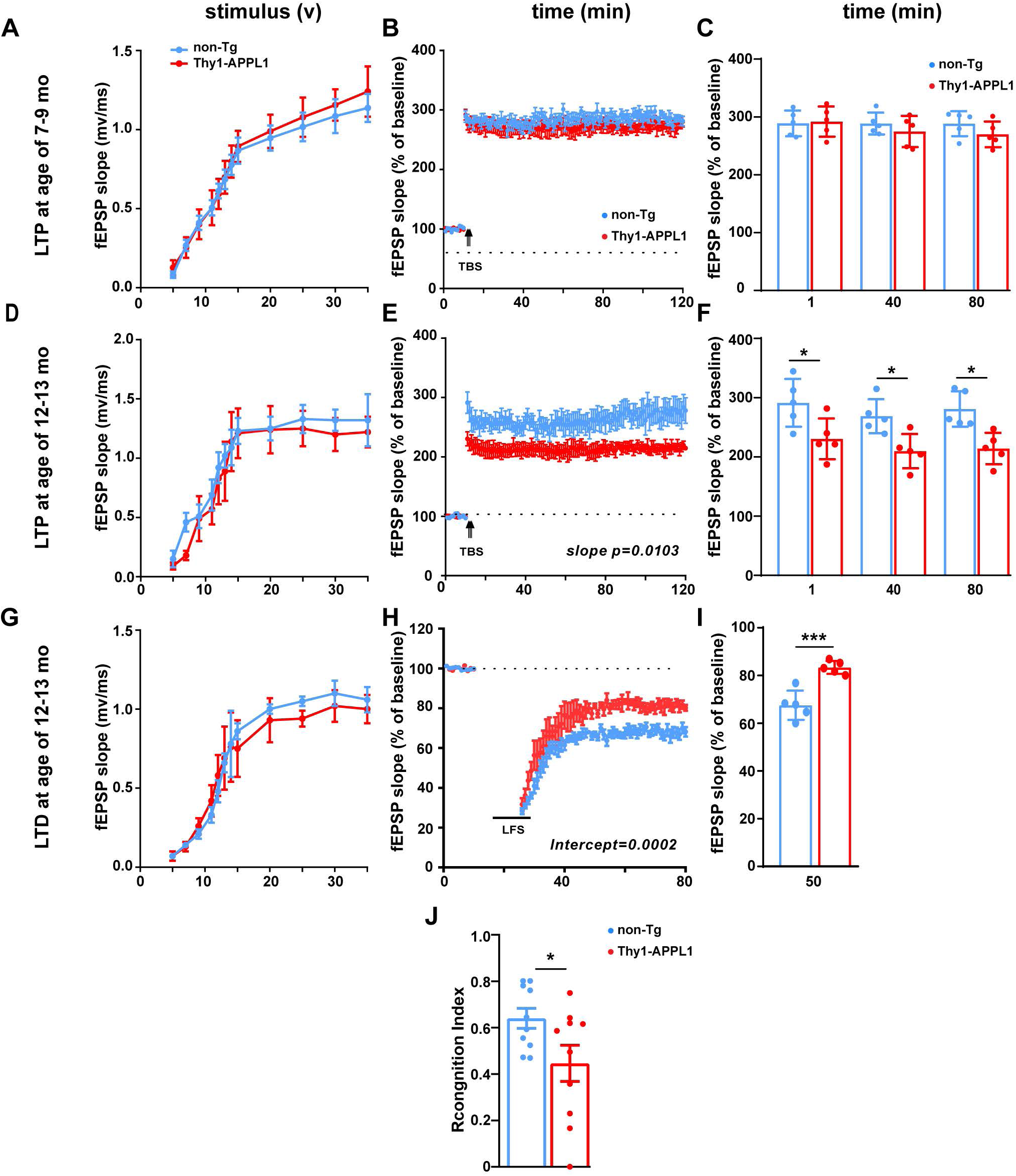
Age-related impairments of synaptic plasticity and hippocampal-dependent memory deficit in Thy1-APPL1 mice. ***A-C.*** non-Tg and Thy1-APPL1 mice at 7-9 months of age (n=5 for both genotype), input/output relationship plots (***A***), LTP by theta-burst stimulation (TBS) in the Schaffer collateral synapses (CA3-CA1) of hippocampal slices (***B)*** and averages of fEPSP slopes at 1, 40, and 80 minutes following tetanic stimulation (***C***) of the hippocampal slices showing no differences between non-Tg and Thy1-APPL1 mice. ***D-F,*** non-Tg and Thy1-APPL1 mice at 12-13 months of age (n=5 for both genotype), input/output relationship plots (***D)***, LTP induced by TBS in the Schaffer collateral synapses (CA3-CA1) in hippocampal slices (***E***) and averages of fEPSP slopes at 1, 40, and 80 minutes (***F***) showing significant reduction in Thy1-APPL1 mice (*F(1, 236)=6.696, p=0.0103* for the slope, linear regression). ***G-I.*** non-Tg, and Thy1-APPL1 mice at 12-13 months of age, (n=5 for both genotypes), input/output relationship plots (***G***), LTD induced by low-frequency stimulation (LFS) (***H***) in hippocampal slices of Thy1-APPL1 (*F(1, 127)=14.83, p=0.0002* for the intercepts, linear regression), and averages of fEPSP slopes at 50 minutes following LFS induction (***I)*** in the hippocampal slices did not show expected reduction in Thy1-APPL1 mice. (***J)*** Recognition index at 3 hours after familiarization indicated the memory deficit in Thy1-APPL1 mice at 12-13 months of age (*t(18)=2.187, p=0.042*). \**p<0.05,* \*\*\**p<0.001,* Two-tailed, unpaired t-test. Data is shown as Mean ± SEM.

## 4. Discussion

Our findings provide strong *in vivo* evidence implicating APPL1 as a key molecular intermediate in a pathogenic cascade propelled by causative and risk genes for AD that lead to abnormal rab5-early endosome signaling and neurodegeneration of basal forebrain cholinergic neurons (BFCN), preceding and predicting the cortical spread of Alzheimer’s pathology (Schmitz and Nathan Spreng, 2016; Schmitz et al., 2018). This cascade is further linked to the decline of memory (Alam and Nixon, 2023), and its detection in prodromal AD (Rozalem Aranha et al., 2023) coincides with the early emergence of signature early endosome enlargement, reflecting rab5 hyper-activation. Clinical trials aimed at pharmacologically attenuating rab5 over-activation have shown encouraging improvement in measures of cholinergic function in Lewy Body Dementia, a disorder in which BFCN loss of function contributes prominently to the clinical presentation (Jiang et al., 2022). The fidelity with which our Thy1-APPL1 replicates the endosomal dysfunction and neurodegenerative consequences seen in AD (Cataldo et al., 1996; Cataldo et al., 2000; Nixon, 2017), APP-based mouse models of AD and DS (Salehi et al., 2006; Jiang et al., 2010; Jiang et al., 2016), and a mouse model of rab5 over-activation (Pensalfini et al., 2020), underscores the important contribution of APPL1 to mediating pathogenic actions of rab5, as suggested by earlier molecular studies of APPL1 in primary neuron cultures (Kim et al., 2016; Xu et al., 2016).

Endogenous APP-βCTF at modestly elevated levels in Down syndrome mouse models enlarges early endosomes, impairing neurotrophic function via retrogradely transported signaling endosomes (Salehi et al., 2006; Xu et al., 2016) and early degeneration of BFCNs (Jiang et al., 2016; Jiang et al., 2022) (Figure 6). Evidence in cell models shows over-expressed APPL1 binds to APP-βCTFs, enlarged early endosomes, and slows axonal transport of endosomes in proportion to the increase in endosome size (Kim et al., 2016). Conversely, in DS fibroblasts, which have enlarged early endosomes due to increased APP-βCTF levels, treating with siRNA to reduce APPL1 levels restores early endosome morphology and function to that seen in diploid cells (Kim et al., 2016). Both APP-βCTF and APPL1 levels are increased in human AD brain (Kim et al., 2016; Johnson et al., 2022), as are the colocalization of APPL1 with rab5-positive endosomes (Kim et al., 2016). Our Thy1-APPL1 mouse directly models an increase in APPL1 expression as reported in human AD, showing that such an increase in APPL1 is sufficient to lead to early endosomal alterations and functional changes in neuron.

**Figure 6.**
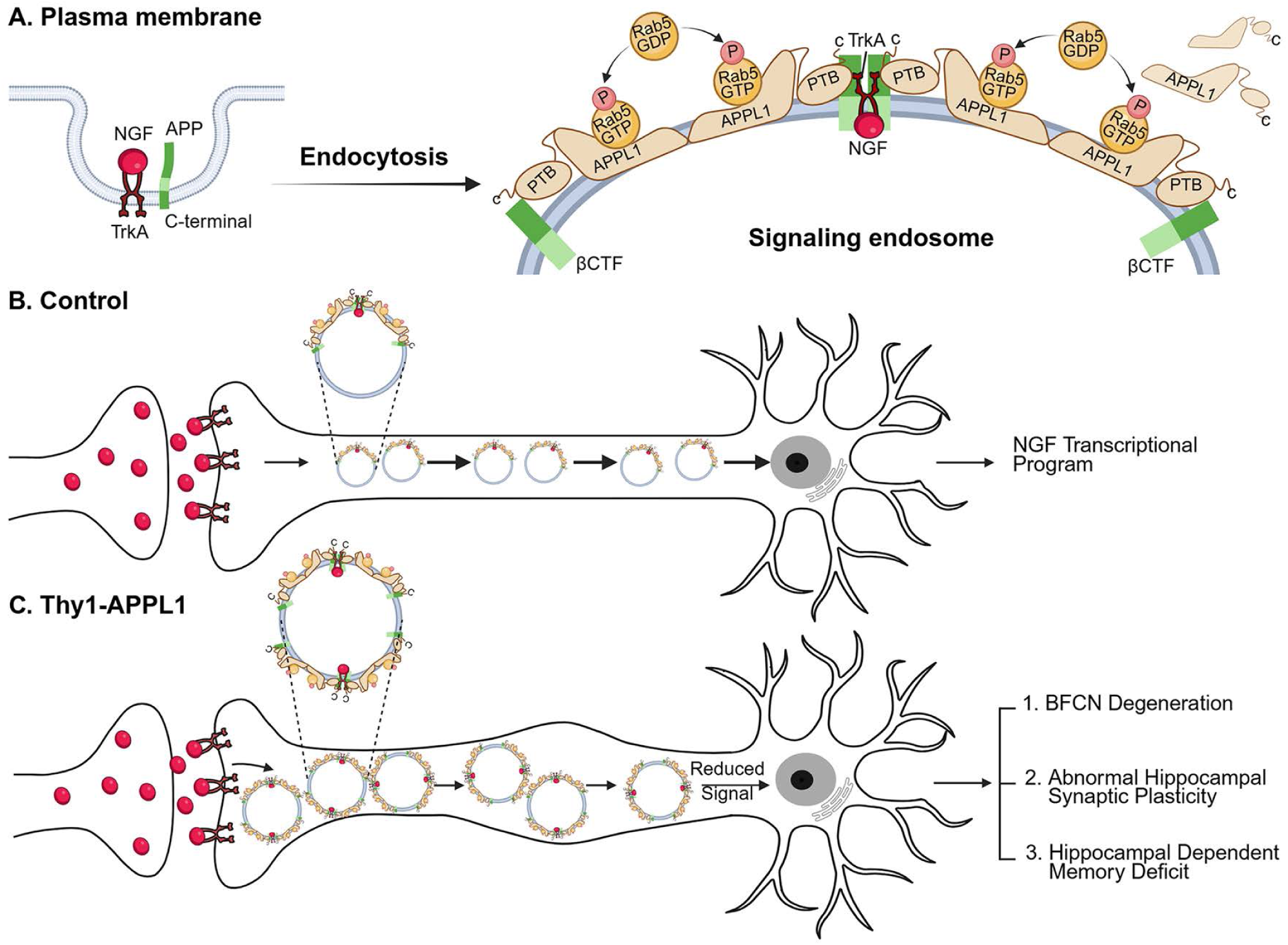
NGF retrograde signalling and involvement of APPL1 in healthy neurons and after a rise in neuronal APPL1 levels related to Alzheimer’s Disease in APPL1 over-expressing neurons. Schematic diagram depicts (***A***) the endocytosis of APP, NGF, and its receptor TrkA into a rab5-early endosome. TrkB mediation of BDNF signalling by APPL1 (not shown) is considered to follow a similar sequence. (***B***) Normal NGF signalling is facilitated by recruitment of APPL1, a direct TrkA ligand and adaptor for other signalling molecules mediating retrograde transport of a maturing endosome carrying the NGF signal to the nucleus to activate a neurotrophic transcriptional program supporting functioning of ChAT neurons and other NGF targets. In AD (not shown-see main text), abnormally elevated APP-βCTF levels arising via multiple possible mechanisms, raise levels of the activated form of rab5 (rab5-GTP) on endosomal membranes, in part by recruiting more APPL1 to the endosome via the phosphotyrosine binding (PTB) domain of APPL1 (Kim et al., 2016). APPL1’s greater affinity for rab5-GTP prolongs association of this activated form on the endosome, thus promoting a pathogenic rab5 hyper-activation leading to increased endocytosis, early endosomal fusion, and endosome enlargement, as depicted in ***C***. **(*C*)** Moderately elevating APPL1 selectively in neurons of Thy1-APPL1 mice phenocopies mouse models of APP-βCTF elevation or rab5 over-expression with respect to rab5 hyper-activation, abnormal endosome enlargement, stasis of endosome transport, synaptic plasticity deficits, and basal forebrain cholinergic neurodegeneration. These pathological effects, as observed in AD brain, also reflect impaired NGF/TrkA signalling and decreased expression of genes for neuronal survival, growth and differentiation (Niewiadomska et al., 2011; Xu et al., 2018). Further information is provided in the text and in more detail in reviews (Nixon, 2017; Nixon and Rubinsztein, 2024).

Given the foregoing *in vitro* evidence in APP over-expression systems and DS fibroblasts (Jiang et al., 2010; Kim et al., 2016), we undertook the current study to further dissect APPL1’s roles *in vivo*. APPL1 recruitment to rab5-endosomes was shown in cell models to be necessary for APP-βCTF-mediated rab5 over-activation, an outcome consistent with APPL1’s action as a stabilizer of the active rab5-GTP complex on endosomes (Kim et al., 2016). With its expression in brain approximately doubled in Thy1-APPL1 mice, APPL1 recapitulated an aging-dependent onset of rab5 over-activation, early endosome swelling (Cataldo et al., 1997; Jiang et al., 2016; Kim et al., 2016; Mathews and Levy, 2019; Pensalfini et al., 2020) and loss of BFCNs, a phenotype invariably seen in AD brain and the various AD models that replicate AD-related early endosome dysfunction (Cataldo et al., 1997; Jiang et al., 2016; Kim et al., 2016; Pensalfini et al., 2020). In the Thy1-APPL1 model, early endosome alterations and these downstream effects were aging-dependent, with early endosomes beginning to enlarge at 7-9 months of age and prior to BFCN loss at 12-13 months of age. Mouse disease models typically show measurable BFCN degeneration months following detectable endosomal alterations (Cataldo et al., 2000; Cataldo et al., 2003; Salehi et al., 2006; Choi et al., 2009; Jiang et al., 2016; Pensalfini et al., 2020; Jiang et al., 2022). Changes in LTP and LTD were also aging-dependent and initially detected at 12-13 months of age, later than the age when early endosome enlargement in cortical neurons was first seen, consistent that the full impact of dysfunctional endosomes plays out over time. That neuronal early endosome enlargement in the Thy1-APPL1 mouse is not seen at 4-5 months of age but detected later is consistent with other models in which the neuronal early endosome appears to be resilient in the younger brain, while this homeostatic ability is lost during aging (Nuriel et al., 2017; Mathews and Levy, 2019). Interestingly, in a model with direct rab5 over-activation (Pensalfini et al., 2020), loss of endosomal homeostasis is apparent early, suggesting that disturbances of rab5 regulatory mechanisms, albeit capable of driving endosomal dysfunction through rab5 over-activation, may be tolerated until the brain begins to age.

NGF signalling is essential for the survival of BFCNs (Sofroniew et al., 1990; Holtzman et al., 1992), and loss of NGF signalling leading to cognitive decline is an early feature in AD and DS (Grothe et al., 2012; Schmitz and Nathan Spreng, 2016). NGF is transduced via endocytosis and the retrograde transport and signalling by a APPL1/rab5 “signalling endosome” containing the NGF receptor TrkA (Xu et al., 2016; Villarroel-Campos et al., 2018). Impaired endosome transport and signalling degrade the NGF trophic stimulus to cholinergic neurons (Salehi et al., 2006; Kim et al., 2016; Kulkarni et al., 2022) (Figure 6). Increasing NGF levels can rescue these neurons (Granholm et al., 2000; Eu et al., 2021), while reducing rab5 over-activation (Xu et al., 2016; Jiang et al., 2022) or APP-βCTF levels (Jiang et al., 2016) prevents endosome dysfunction and preserves cholinergic function. Our study therefore offers the possibility of targeting rab5 via modulation of APPL1 to reactivate functional cholinergic neurons and reverse disease progression.

Increased phosphorylated tau, as detected by PHF1, was also measured in synaptosomes isolated from the Thy1-APPL1 mice (Fig. 3). Endosomal pathway alterations are known to impact tau phosphorylation as well as the spread and seeding of pathogenic tau within the brain (Knopman et al., 2021). APPL1 influence on multiple protein kinases involved in axonal transport (for example, Akt, GSK3ß, MAP Kinase) (Mitsuuchi et al., 1999; Diggins and Webb, 2017) is also likely to affect tau phosphorylation directly in the Thy1-APPL1 model.

Rab5 is associated with both pre- and post-synaptic vesicles, where rab5 is critical in mediating function by regulating synaptic vesicle endocytosis and thus membrane and protein cargo retrieval (Star et al., 2005; Chanaday et al., 2019). These are also endosomal functions vulnerable in neurodegenerative diseases (Bonnycastle et al., 2021; Harper and Smillie, 2021; Bellucci et al., 2022). Synaptosomes containing numerous synaptic vesicles have been used as an *ex vivo* system to study synaptic dysfunctions in AD and other neurodegenerative diseases (Evans, 2015; Jhou and Tai, 2017; Ahmad and Liu, 2020; Griffiths and Grant, 2023). We observed that increased APPL1 expression induces synaptosome endocytosis, consistent with its activating rab5 on neuronal endosomes. Both knockdown mediated by siRNA and an APPL1 functional knockdown mediated by inhibitory peptides *in vitro* show that APPL1 is necessary for normal hippocampal LTP and LTD (Fernandez-Monreal et al., 2016; Wang et al., 2016; Wu et al., 2020; Formolo et al., 2022; Hua et al., 2023). We find that APPL1 overexpression *in vivo* also impacts hippocampal electrophysiology. Given APPL1’s functions at the early endosome, both under- and over-expression of APPL1 are likely to perturb early endosome signaling as well as the formation and cargo-acquisition of early endosomes through endocytosis (Diggins and Webb, 2017). Overexpression of APPL1 in the Thy1-APPL1 mouse increased rab5 activity, driving synaptic endocytosis. It is also likely to lead to entrapment of endocytic cargo, which enlarges early endosomes preventing appropriate synaptic vesicle recycling and trafficking. The increase of APP-βCTFs seen in the synaptosomes of Thy1-APPL1 mouse is also consistent with endocytosed cargo being entrapped in early endosomes. BACE1 cleavage of APP to generate APP-βCTFs occurs in endocytic compartments, and increasing the uptake and dwell-time of APP within endosomes increases APP-βCTF generation (De Strooper and Karran, 2016). Abundant APPL1 at endosomes, given its interaction with endosomal APP-βCTFs, is likely to increase the dwell time of the rab5/APPL1/APP-βCTF complex, further accentuating rab5 activation.

Among its various neuronal signaling functions (Lin et al., 2006; Deepa and Dong, 2009; Lee et al., 2011; Hupalowska et al., 2012), APPL1 acts as a molecular link between APP-βCTF and rab5 (Kim et al., 2016), a critical cross-roads for pathways that converge on the early endosome during AD pathobiology. The Thy1-APPL1 mouse highlights the important role APPL1 can play in transducing APP-βCTF-signaling in AD to rab5, thus regulating early endosome function, growth factor signaling, and cholinergic neuronal functioning. We now show that overexpression of APPL1 leads to phenotypes similar to those that result from the direct over-activation of rab5 (Pensalfini et al., 2020). Whether other risk factors for AD-driven early endosome and synaptic dysfunction, such as APOE4, BIN1, CD2AP, PICALM, RIN3, SORL1 and others (Nixon, 2017; Perdigão et al., 2020) act through APPL1 is unknown; nevertheless, the Thy1-APPL1 model further demonstrates that multiple molecular mechanisms leading to AD-like early endosome dysfunction are known to converge on rab5 and its regulation.

## Author contributions

R.A.N. and Y.J. designed the research; Y.J., K.S., M.J.B., S.S., B.S.B., C.B., J.F.S., C.N.G., M.P., P.H.S., J.P., A.P., S.M. and L.W. conducted the research; Y.J. wrote the first draft; Y.J., P.M.M., and R.A.N. edited the paper.

## Acknowledgments

The study was supported by NIH grants P01AG017617 and R01AG062376 grants to R.A.N. B.S.B. and S.S. are supported by NIH grant R01AA29686. P.M.M is supported by RF1AG088226. We gratefully acknowledge the excellent assistance of Swati Jain in the preparation of this manuscript for submission and for the illustration of Figure 6.

## Abbreviations

AD: Alzheimer’s disease
APP: amyloid precursor protein
APPL1: adaptor protein containing pleckstrin homology domain, phosphotyrosine binding domain and leucine zipper motif
BFCN: Basal forebrain cholinergic neurons
ChAT: choline acetyltransferase
CMFDA: 5-chloromethyl fluorescein diacetate
DAB: diaminobenzidine
DS: Down Syndrome
EM: electron microscopy
fEPSP: field excitatory postsynaptic potential
FM^TM^4-64: F4-64: (*N*-(3-Triethylammoniumpropyl)-4-(6-(4-(Diethylamino) Phenyl) Hexatrienyl)] Pyridinium Dibromide)
HRP: Horseradish peroxidase
LTD: long-term depression
LTP: long-term potentiation
MSN: medial septal nucleus
NGF: Nerve growth factor
NOR: Novel Object Recognition
PHF1: paired-helical filament tau, phosphorylated at Ser396 and Ser404 site,
RI: Recognition Index
SVE: synaptic vesicle endocytosis.

